# A model for within- and between-host evolution in pathogens

**DOI:** 10.64898/2026.09.24.754225

**Authors:** James Crescenzi, Alex McAvoy, Parul Johri

**Author notes:** Correspondence may be sent to these authors;. These authors contributed equally to this work (Alex McAvoy, Parul Johri).

## Abstract

Many pathogenic and microbial species undergo complex life cycles wherein they experience drasrtic bottlenecks during invasion into a host and rapid growth within hosts. The effects of such a complex population history on patterns of genetic variation is not currently understood. We model a pathogen population as a metapopulation where each deme represents a host and incorporate transmission dynamics as well as the effects of transmission bottlenecks. We employ a coalescent framework, obtaining analytical expressions for pairwise times to coalescence and genetic differentiation within and between hosts. We find that recurrent bottlenecks rescale the coalescent process within hosts to the size of the bottleneck, reducing within-host variation. In addition, the pairwise distribution of time to coalescence under our model deviates substantially from the Wright-Fisher process when the number of hosts (*d*) is of similar order to the size of the bottleneck (*k*), i.e., *k* ∼ *d* and when 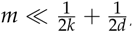, where *m* is the migration rate. Otherwise, the pairwise process under our model converges in distribution to the Wright-Fisher process. Coalescent simulations show that under the same conditions the site frequency spectrum (SFS) under our model deviates from that under the Kingman coalescent and is drastically skewed towards singletons. Our results suggest that the inference of selection or demographic history from pathogen population genetics data may be systematically biased by the mechanics of pathogen reproduction, indicating a need for further theoretical work that can inform inference.

## Introduction

As whole genome sequences of multiple human pathogens have become available, both within and across species, it is becoming more feasible to employ population genetic approaches to uncover their selective and population history. We now have more than 17 million genomes of SARS-CoV-2 available (Smith et al. 2026) and variants from at least 30,000 genomes of *P. falciparum* (MalariaGEN 2025), the primary causative agent of malaria. However, a major challenge in the interpretation and application of population genetics approaches to human pathogens is the complex demography experienced by most pathogens. Because pathogens continuously invade new hosts, they undergo drastic bottlenecks during invasion and exponential growths within hosts. The effect of such a complex population history on patterns of genetic variation is not currently understood.

Multiple organisms have been shown to experience recurrent generational bottlenecks (Table 1). For instance, *P. falciparum* has a within-host carrying capacity of approximately 10^9^ − 10^13^ individuals, but each generation bottlenecks down to 10-100 parasites in a human host (Graumans et al. 2020; Henry et al. 2025). SARS-CoV-2 has a carrying capacity of 10^9^-10^11^ parasites (Sender et al. 2021) and an effective bottleneck size of 1-8 individuals (Lythgoe et al. 2021), while influenza A has a carrying capacity of at least 3.28 × 10^8^ (To et al. 2010) and an effective bottleneck size of 1.68 (McCrone et al. 2018) with 1-13 census individuals (Ghafari et al. 2020). Importantly, organelles like mitochondria share a similarly complex life cycle with repeated expansions within hosts followed by severe bottlenecks. A strong germline bottleneck occurs in mitochondria (reviewed in Zhang et al. 2018) with an estimated effective size of 10-30 mitochondrial DNA segregating units in humans (Zaidi et al. 2019), corresponding to ∼ 1,500 mitochondrial genomes in humans (Floros et al. 2018) and 200-1,000 in mice (Cao et al. 2007). Moreover, pests (e.g., bed bugs; Fountain et al. 2014) and microbial organisms with symbiotic (e.g., *Buchnera*; Funk et al. 2001) or parasitic lifestyles (e.g., *Wolbachia*; Pietri et al. 2016), including species comprising the microbiome, may experience similar recurrent bottlenecks when invading new hosts. Although a few previous studies have emphasized the importance of studying within-host processes and modeled the effects of transmission bottlenecks (e.g., Roze et al. 2005; Kennedy and Dwyer 2018) on specific evolutionary parameters of interest, a common framework is missing. There is thus a need to build an evolutionary null model accounting for within-host diversity and repeated bottlenecks for microbial organisms and human pathogens.

**Table 1:**
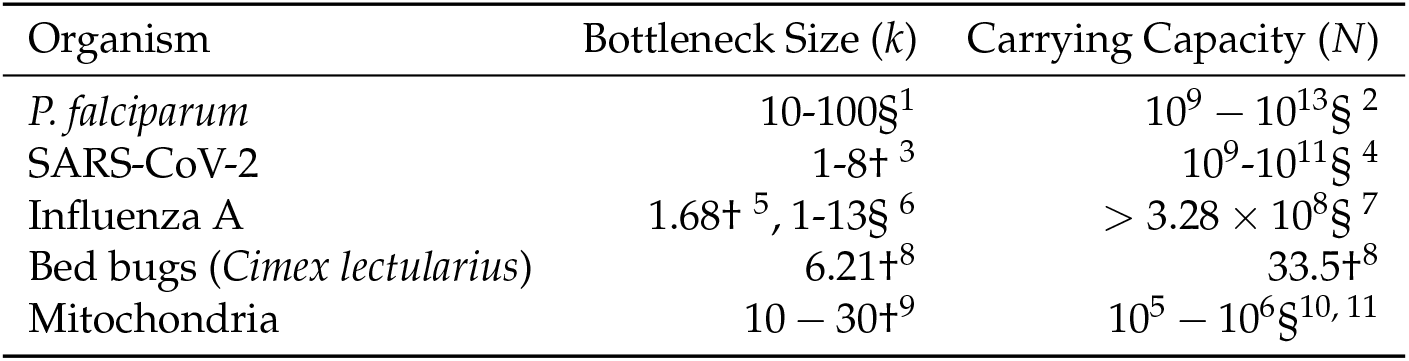
Empirically estimated values of bottleneck size (*k*) and carrying capacity (*N*) from previous studies. Estimates of census size are denoted by §, while estimates of an effective size are denoted by †. 1, 2: Kappe et al. 2010, Graumans et al. 2020, Henry et al. 2025, 3: Lythgoe et al. 2021, 4: Sender et al. 2021, 5: McCrone et al. 2018, 6: Ghafari et al. 2020, 7: To et al. 2010, 8: Fountain et al. 2014, 9: Fountain et al. 2014; Zaidi et al. 2019, 10: Floros et al. 2018, 11: Craven et al. 2010

We employ a metapopulation framework to account for some of the peculiarities of pathogen reproduction. Metapopulations, introduced by Sewall Wright (1931) have been historically well studied in population genetics. In particular, the properties of Wright’s island model, where each deme is equally likely to receive migrants from another deme, have been studied extensively (Moran 1959). For instance, Maruyama (1970) and Latter (1973) described the properties of identity by descent and the effective number of alleles in Wright’s island model. Wakeley (1998) considered properties of the coalescent generated by the island model in the infinite-demes limit, using simulations to study the site frequency spectrum, and Nagylaki (1998) studied the conservative, weak, and strong migration limits. Metapopulation structure was also shown to affect the site frequency spectrum (Wakeley and Aliacar 2001) and Tajima’s *D* (Tajima 1989; Pannell 2003).

Here, we develop a general mathematical framework, using the coalescent, to study evolutionary dynamics in organisms that experience repeated bottlenecks. We propose a metapopulation framework where each deme represents a host, transmission every generation incorporates a bottleneck, and migration between demes represents mixed infections. We use a first-step analysis to describe pairwise diversity and time to coalescence within and between hosts, and examine their distributions. We use simulations to describe the site frequency spectrum and when it deviates from the SFS under standard models (the Kingman coalescent).

## Model

We describe pathogen evolution as a metapopulation process, in which there are *d* demes (representing hosts), each of fixed size, *N*. Time proceeds in discrete, non-overlapping generations. Each deme *i* ∈ {1, . . ., *d*} independently samples a parental deme uniformly at random with replacement. Parents may be sampled more than once or not sampled at all, leading to stochasticity in the transmission process. We refer to this as *deme drift*. The infection cycle begins with a transmission bottleneck, where the number of pathogens within a single host grows rapidly from *k* to *N*, where *N* represents the host’s carrying capacity and *k* ≪ *N* (typically). We model this behavior by considering an unobservable intermediate sampling step between deme *i* in generation *t* + 1 and its parent. A propagule of size *k*, representing the few pathogens that make it through the transmission bottleneck, is sampled from the parent with replacement. Then, *N* individuals are sampled from the propagule to form deme *i*. We assume that the propagule expands instantaneously to size *N* within the deme, with each of these *N* individuals choosing a parent from the propagule pool uniformly at random, with replacement.

For each of the *k* founders of deme *i* at generation *t* + 1, we sample a parent from the parental deme with probability 1 − *m*, and sample a parent from the migrant source deme with probability *m*. The *k* founders expand to *N* individuals via uniform sampling with replacement. Note that generation *t* + 1 starts when the *k* founders expand to *N*. This modeling choice was made in order to avoid pairs of individuals sampled in generation *t* coalescing in generation *t*. Because the intermediate sampling step is unobservable, individuals that first share an ancestor during the intermediate step between generations *t* and *t* + 1 will appear to coalesce during generation *t* + 1. See a visual depiction of this model in Figure 1.

**Figure 1:**
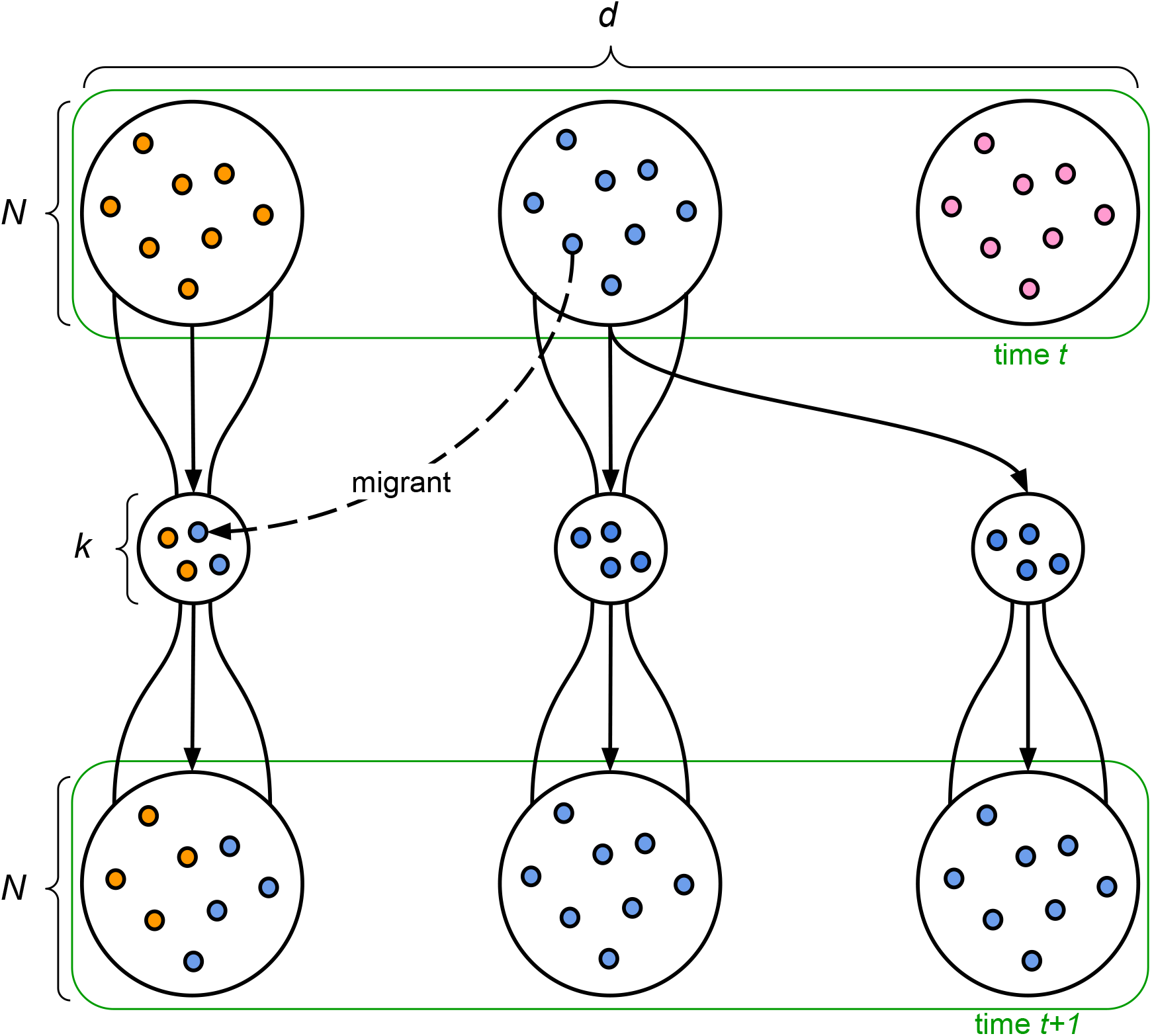
A visual representation of our model capturing pathogen dynamics in a population. Small solid colored circles represent pathogen/microbial genomes with different colors representing different haplotypes. Larger circles represent hosts (demes). Migration represents a coinfection. Here *d* is the number of demes, *N* is the number of pathogen individuals within hosts, *k* is the number of pathogen individuals during the bottleneck, and *t* is time in generations.

We model coinfection (i.e., a single host being simultaneously infected by two different parental hosts) as migration, where each of the *k* individuals in the propagule have probability *m* of being a migrant. We assume that each deme samples a migrant source deme uniformly from the other *d* − 1 demes and that all migrants in the propagule come from the same migrant source deme. This assumption is realistic for many vector-borne diseases like malaria, as well as pathogens with contact-based transmission such as Influenza A and SARS-CoV-2. However, there may be organisms where drawing migrants from the metapopulation as a whole would be more appropriate, such as waterborne diseases caused by *Vibrio cholerae* or *Campylobacter jejuni*, which we do not focus on here. We define one generation to be one full infection cycle, i.e., the time between invading the first host to the next one.

## Results

### Genetic differentiation between hosts

We use a coalescent approach to study the patterns of genetic diversity produced by our model of pathogen reproduction. We are interested in time to coalescence for a pair of individuals in the same deme (*T*_0_), different demes (*T*_1_), and sampled from the whole metapopulation (*T_T_*). We use these times to coalescence to calculate expected nucleotide diversity within demes (*π_S_*), between demes (*π_B_*), and from the whole metapopulation (*π_T_*). In our model, nucleotide diversity is a random variable obtained by sampling the coalescence time for the two individuals in question, and then counting the total number of mutations along the two diverging branches, where at each generation a mutation happens independently with probability *u*. Throughout, we assume that *k*, *d*, *m*, *N*, and *u* are fixed and do not change over time.

### Mean time to coalescence

To obtain the distribution of times to coalescence of any two individuals under our model, we describe the stochastic behavior of pairs going backward in time, using an absorbing Markov chain. A pair of individuals can be in one of three states: in the same deme (state 0), in different demes (state 1), and coalesced (state 2). The transition matrix between states, *P*, can be partitioned into blocks,

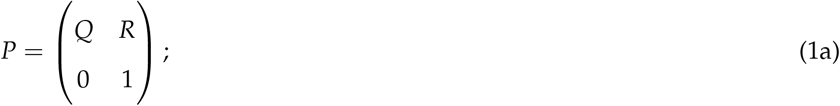

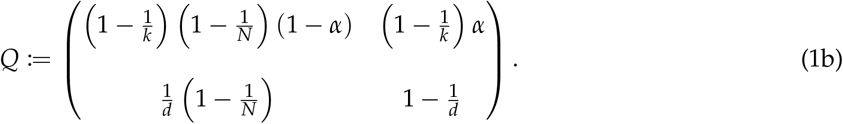

*Q* gives the probabilities of transitioning among transient (non-coalesced) states, and *R* represents the probability of coalescing from each transient state. Since there is only one absorbing state, *R* is uniquely determined by *Q* as *R_i_* = 1 − *Q_i_*_,1_ − *Q_i_*_,2_. Note that *α* = 2*m* (1 − *m*) represents the probability that a pair of individuals in the same deme have parents in different demes due to migration.

The probability of being in state *j* after *t* steps, given that the process starts in state *i*, is P (*X_t_*= *j* | *X*_0_ = *i*) = (*P^t^*)*_i_*_,*j*_. The 2 × 2 submatrix in the upper-left part of *P^t^* is equal to *Q^t^*, which lets us compute the probability that a pair of lineages has not yet coalesced as P (*T > t* | *X*_0_ = *i*) = (*Q^t^*)*_i_*_,1_ + (*Q^t^*) *_i_*_,2_ = (*Q^t^***1**)*_i_*. Rearranging, we see that P (*T* ⩽ *t* | *X*_0_ = *i*) = [*I* − *Q^t^* **1**]*_i_*, where **1** is a vector of ones, which gives

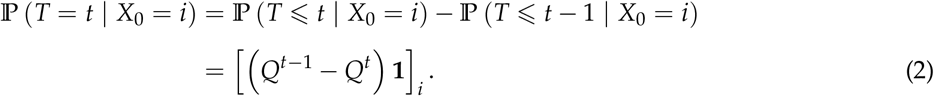

This expression immediately yields equations for the distributions of *T*_0_, *T*_1_, and *T_T_*, since P (*T*_0_ = *t*) = P (*T* = *t* | *X*_0_ = 0), P (*T*_1_ = *t*) = P (*T* = *t* | *X*_0_ = 1), and 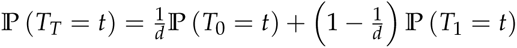.

Since *Q* is substochastic, the number of steps to absorption is finite with probability one. Letting *F* be the fundamental matrix *F* = (*I* − *Q*)^−1^, the vector of conditional expected times to coalescence is given by *e* = *F***1**, and E[*T_i_*] = *e_i_*. The unconditional expectation is given by E [*T_T_*] = *µF***1** = *µe*. The vector of conditional variances of time to coalescence is *V* = (2*F* − *I*)*e* − *e_sq_*where *e_sq_* is the elementwise product of *e* with itself. *Var*(*T_i_*) = *V_i_*, and the unconditional variance of time to coalescence is *Var*(*T*) = *µV* + *µe_sq_*− (*µe*)^2^. While explicit equations for the variance are somewhat complicated, the expected coalescence times simplify to

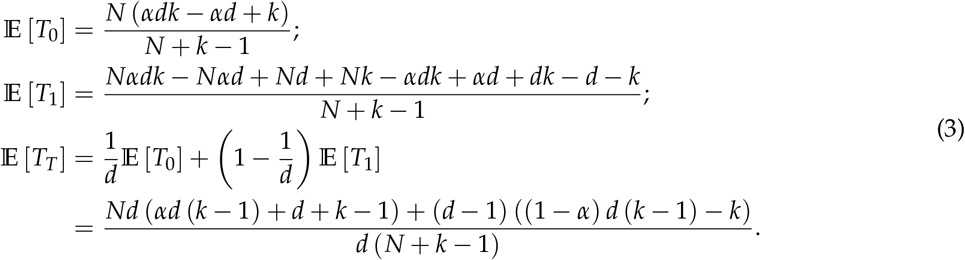

When hosts carry large numbers of pathogens, taking *N* → ∞ gives

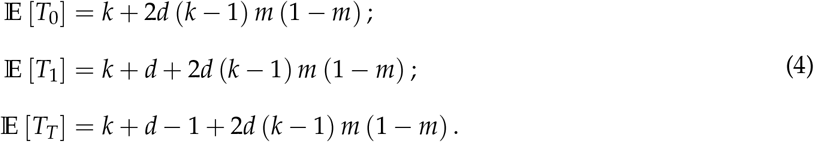

It is noteworthy that, as *N* → ∞, E [*T*_1_] − E [*T*_0_] = *d*, E [*T_T_*] − E [*T*_0_] = *d* − 1, and E [*T*_1_] − E [*T_T_*] = 1.

The process of coalescence under our model has a few phases: *(i)* pairs in different demes are collected (Wakeley and Aliacar 2001) into the same deme. This process has expected time *d* for pairs of individuals in different demes, and is not relevant for pairs in the same deme. It is also independent of migration, because deme drift has the same probability of placing individuals into the same deme if one, both, or neither of the individuals are migrants. *(ii)* Pairs of individuals that are in the same deme coalesce with expected time *k*, *or (iii)* are separated into different demes by migration. Then, time to coalescence is the sum of these three random processes: collecting lineages in a deme (unless they already share a deme), coalescence, and separation that repeats the process. In the large-*N* limit, the expected time for the collecting process to occur when lineages start in different demes is *d* generations.

### Nucleotide diversity

The expectation of *π_S_*, *π_B_*, and *π_T_* can be found by computing the expectation of a binomial random variable with rate *u* (representing the neutral mutation rate per site per generation) and 2*T* trials. E[*π_S_*] = 2*u*E [*T*_0_], E[*π_B_*] = 2*u*E [*T*_1_], and E[*π_T_*] = 2*u*E [*T_T_*]. The variance in *π* can be computed similarly as

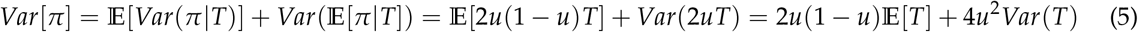

To confirm our analytical expressions of pairwise summary statistics, we simulated the coalescent process for a pair of individuals that have the same ancestral process as our model. The pairwise simulations are described in the supplement (Text S1). The expectation and variance of *π* are calculated across 200,000 replicates.

In Figure 2, we plot expected nucleotide diversity and *F_ST_*, which measures the proportion of genetic variation attributable to differences between populations, across a range of migration rates. *F_ST_* is computed numerically from our simulations as 1 – *π_S_*/*π_T_*, where *π* is the mean *π* for a (*k*, *m*) pair. Observe that for low values of *m*, the population behaves as though *m* is zero. There is an increase in diversity at an intermediate value of *m* and for values of *m* greater than 0.5, there is a decrease in *π*, as the model behaves symmetrically around *m* = 0.5 due to how we model migration. When migration is sufficiently uncommon, pairs of lineages coalesce as though migration does not happen; this is the case when migration almost never occurs in the history of two lineages. We quantify this based on the expected time to coalescence of any pair of individuals, E [*T_T_*]. Assuming that *m* is small enough that *m*^2^ is negligible, we have E [*T_T_*] ≈ *k* + *d* − 1 + 2*d* (*k* − 1) *m*, so from the perspective of E [*T_T_*], migration is “rare” whenever *m* ≪ (*k* + *d* − 1) / (2*d* (*k* − 1)). In this regime, *π_S_* scales linearly with *k* and *π_T_* scales linearly with *k* + *d*. Thus while within-host diversity is drastically reduced by bottlenecks and is entirely determined by *k*, diversity across the entire metapopulation is only slightly affected by the bottleneck size, especially if *d* is much larger than *k*.

**Figure 2:**
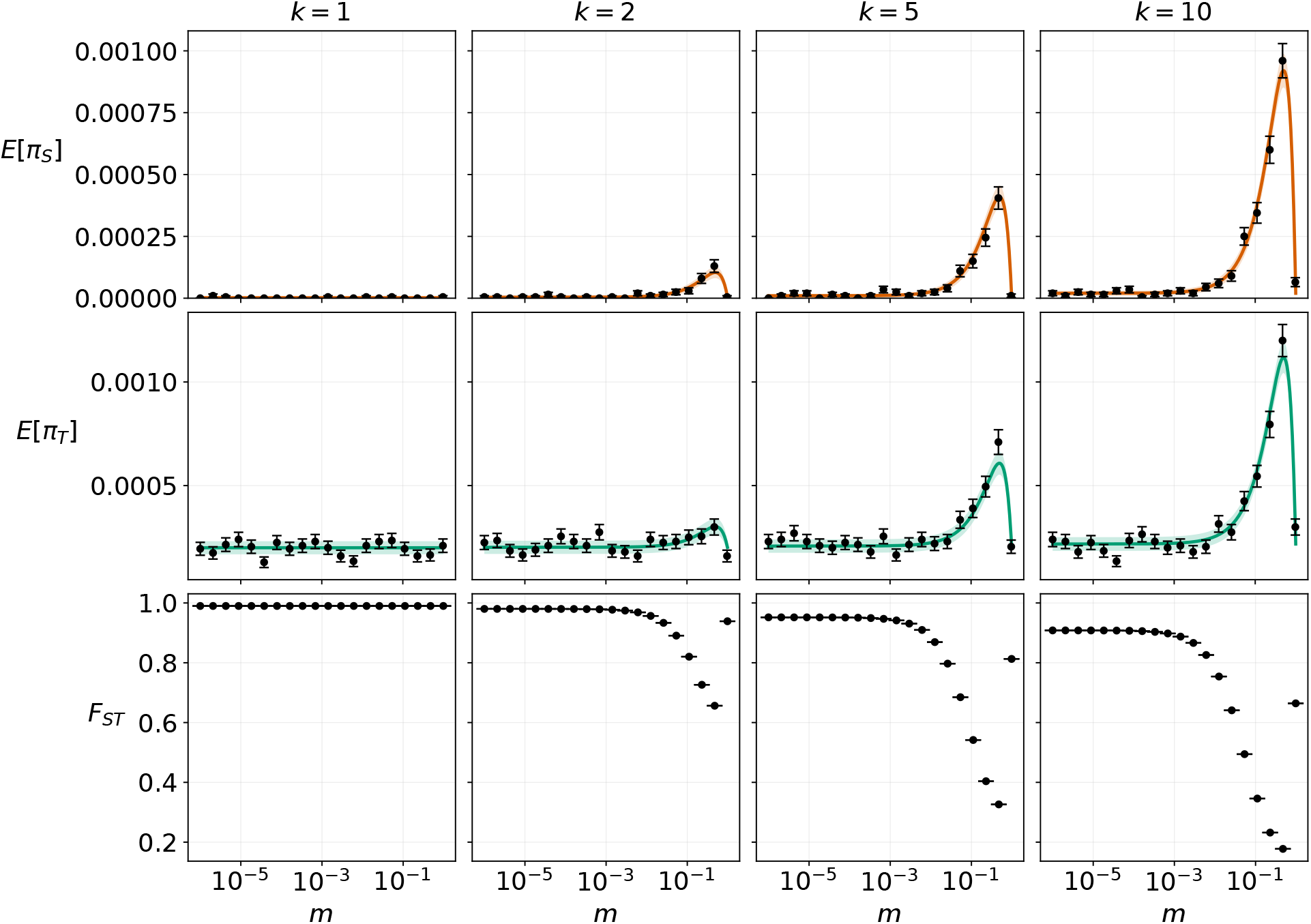
Comparison of theoretically calculated *π_S_* and *π_T_* with results from 200,000 (= *n_r_*) replicates of coalescent simulations for varying bottleneck sizes (*k*) and migration rates (*m*). Here *u* = 10^−6^, *d* = 100, and *N* = 10^10^. The theoretical means of each *π* statistic are given by the lines, while the ribbons represent the expectation of the standard error 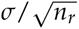. The simulated means are given by the points, while the error bars represent the standard error. As *n_r_* → ∞ the standard errors approach zero, and the simulated means match the theoretical expectations exactly. This is demonstrated by Figure S1. We compute *F_ST_*empirically from each simulation replicate and do not estimate it theoretically.

### The distribution of time to coalescence

We obtain the distribution of coalescence times under our model (Figure 3). Because the analysis of population genetics data most often assumes a Wright-Fisher population, we contrast the distribution of *T_T_* against the distribution of time to coalescence under a Wright-Fisher model (*T*_WF_) with *N_e_*= E [*T_T_*]. While another metapopulation model, such as Wright’s island model, may seem like a good choice for comparison, it is not straight-forward. Unlike in our model, the collecting process for the classical metapopulation models depends on migration. Under the island model, for the collecting and coalescing processes to be of similar duration requires that either *(i)* migration must be sufficiently common that 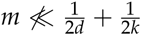 or *(ii)* the probability of coalescence within a deme must be small enough to be of order *m* when *m* is small. The first case would require the full distribution of times to coalescence across *m* without invoking an infinite ambient population limit. The second case is incompatible with our assumption of generational bottlenecks.

**Figure 3:**
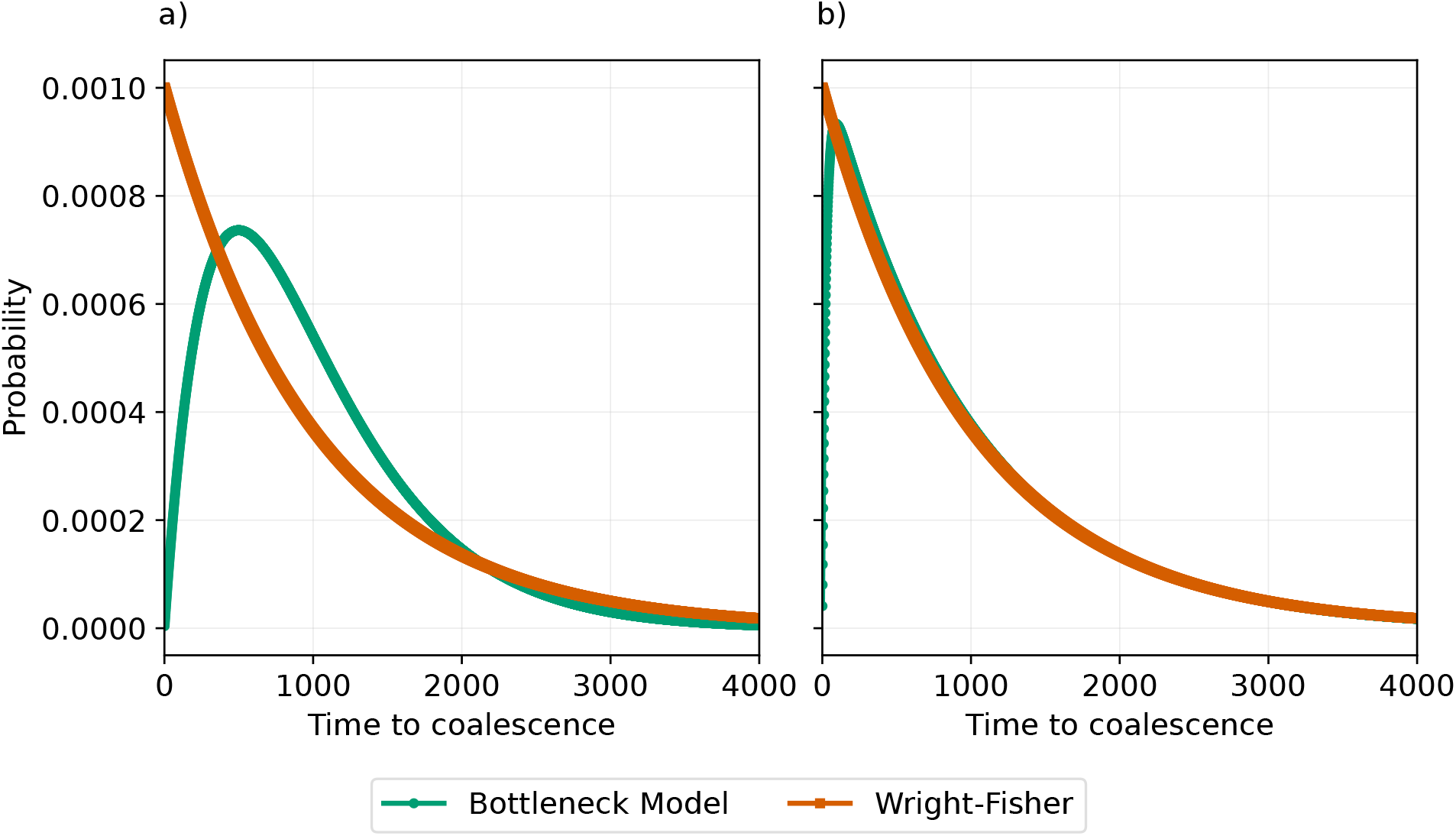
Comparison of the distribution of time to coalescence in our model (*T_T_*) with the distribution of time to coalescence in a Wright-Fisher model with *N_e_*= E [*T_T_*]. In (a): d = 500 and k = 500. In (b): *d* = 975, *k* = 25. In both panels, *m* = 0 and *N* = 10^10^, consistent with the *N* limit. Demonstrates that the distributions can be very different when *k* ∼ *d*, but converge when *k* ≪ *d* or *k* ≫ *d*.

Time to coalescence under the Wright-Fisher is geometrically distributed: its most likely value of *t* is 1, and its distribution function decreases monotonically. Conversely, under our model coalescence follows a phase-type distribution that has low probability for low *t*, and peaks with non-trivial *t*. We see from Figure 3 that these two distributions can be both different and similar depending on the parameters used. Figure 3a gives an example of when these distributions are most different: when *k* and *d* are of the same order of magnitude and *m* is small. Figure 3b gives an example of how the distributions become more similar when *k* and *d* are of different orders of magnitude: the peak of the distribution of *T_T_* is pushed toward zero, causing it to take a similar shape to the distribution of *T*_WF_.

In order to systematically investigate the relationship between *T_T_* and *T*_WF_, we calculate the total variation distance,

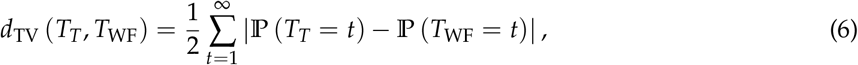

and plot it across a range of *k* and *d* values (Figure 4). The migration rate is such that 2*kdm* = *k* + *d* when *k* = *d* = 5,000. When *k* and *d* are small enough that 2*kdm* ≪ *k* + *d*, the distributions *T_T_* and *T*_WF_ are most different when *k* = *d* and the TV-distance decreases symmetrically when either one of *k* or *d* dominates the other (i.e., the TV surface forms a ridge when *m* is small and *k* = *d*). As *k* and *d* increase, they become large enough that 2*dkm* ≪̸ *k* + *d* and the TV-distance falls off accordingly (i.e., the ridge decreases in magnitude).

**Figure 4:**
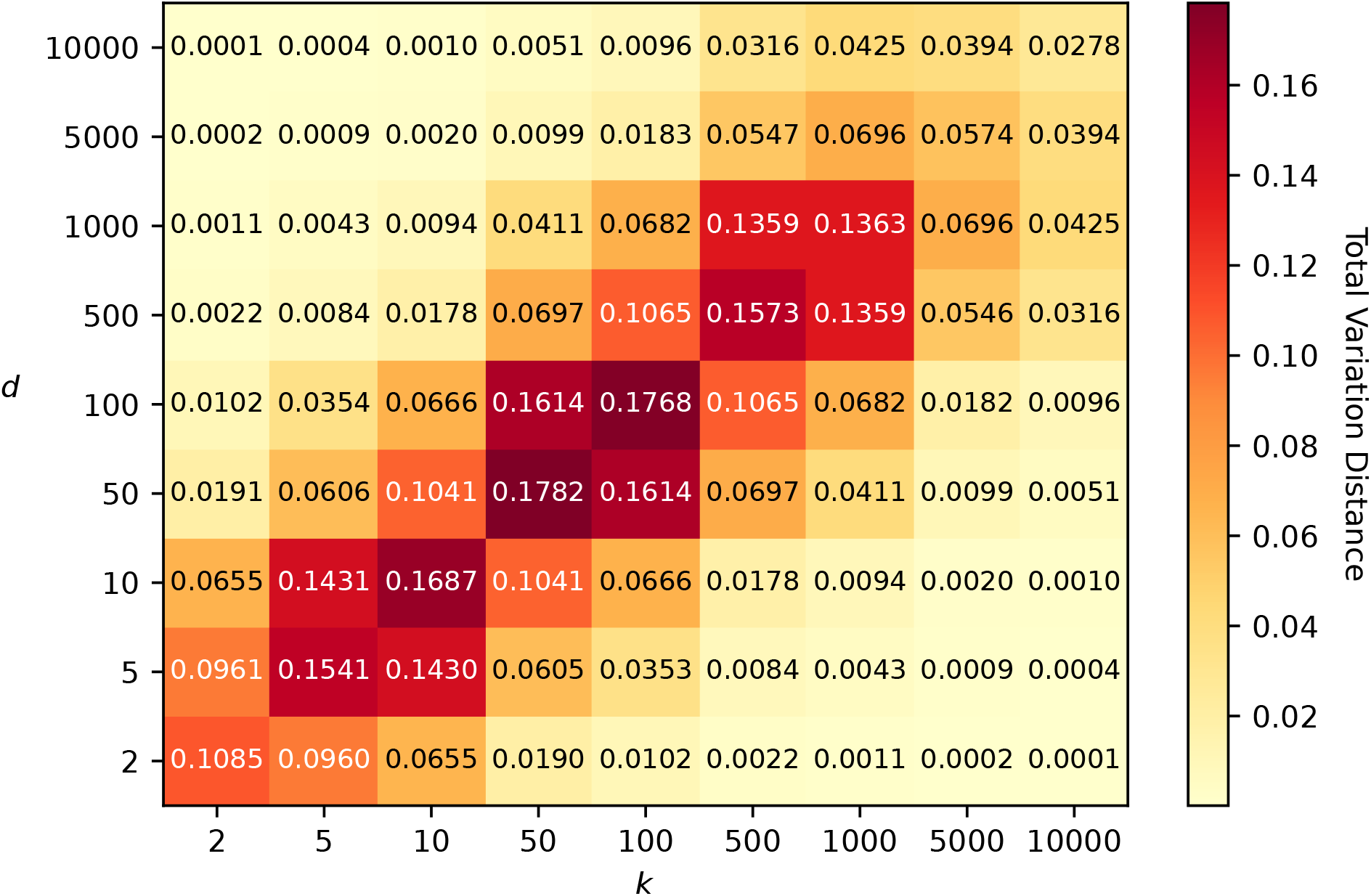
The total variation distance calculated between the distribution of *T_T_* and time to coalescence under a Wright-Fisher model with *N_e_* = E [*T_T_*]. The migration rate *m* = 0.0002 (2*dkm* = *k* + *d* when *k* = *d* = 5,000), and *N* = 10^10^.

We can understand this behavior by considering a separation of timescales. When *k* is much less than *d*, lineages that start in different demes coalesce in approximately *d* generations; coalescence within a deme occurs almost immediately compared to the collecting process. Conversely, when *d* is much less than *k*, lineages in different demes are placed into the same deme by deme drift almost immediately; lineages that start in different demes coalesce in about *k* generations. It is only when *k* and *d* are of similar orders of magnitude that the collecting and coalescing processes can interact with each other, yielding the phase-type distribution our model generates. When the migration rate *m* is large (and *k* is not trivial), lineages that enter the same deme either coalesce immediately or are separated by migration. The process of coalescence therefore becomes waiting for a pair of lineages to enter the same deme *and* coalesce immediately, which is roughly geometrically distributed.

### The site frequency spectrum

Because bottlenecks can dramatically alter the distribution of allele frequencies, we asked whether the site frequency spectrum (SFS) of derived alleles in the metapopulation was significantly different from that generated by the Kingman coalescent with *N_e_* = E [*T_T_*] (Kingman 1982; Fu 1995). This is reasonable because the SFS in an island model with a large number of demes also converges to that in a Wright-Fisher population (Wakeley 1998). Moreover, under the same infinite ambient population assumptions under which the Wright-Fisher model generates the Kingman, our model generates the Kingman when *m* = 0 under the limit *N* = *k*; *N*, *k* → ∞. This can be shown by demonstrating that the collecting process occurs instantaneously when measuring time in units of *N*/2, and that the discrete-time transition probabilities of any two states in the coalescent process can be rescaled by *N*/2 so that the resulting continuous time process is the Kingman coalescent. A similar separation of timescales argument could be used to show that the Kingman coalescent is generated with small *k*, *N* in the limit as *d* → ∞.

We want to establish an informal link between deviations in the distribution of pairwise time to coalescence under our model from the Wright-Fisher model and deviations in the distribution of allele frequencies under our model from the Kingman coalescent, which arises from the Wright-Fisher model (Kingman 1982). Because the SFS is influenced by the allele-age frequency (Griffiths 2003), which is in turn affected by the pairwise probability of coalescence, it is reasonable to suggest that the SFS will deviate from the Kingman when the TV-distance between *T_T_* and *T*_WF_ is large. The natural question becomes: how does the SFS under our model deviate from the Kingman, and what values of the TV-distance correspond to significant deviation in the SFS?

The pairwise TV-distance is largest when *k* and *d* are equal and *m* is small (Figure 4). We therefore study how the SFS under our model deviates from the Kingman when *m* = 0.0002 (which gives 2*dkm* = *k* + *d* when *k* = *d* = 5,000) and *k* and *d* vary (Figure 5). When *k* and *d* are equal, the largest TV-distance of approximately 0.0574 is observed (Figure 4). The deviation in the SFS is massive; there are more than twice as many singletons under our model as there are under the Kingman SFS (Figure 5c). Even when *k* and *d* differ by one order of magnitude (*k* = 500, *d* = 9,500, TV-distance 0.033; Figure 5b), we observe a large deviation in the SFS. Under our model, we see about 1.6 times the number of singletons as in the Kingman. The same holds when *k* is greater than *d*, and with *k* = 9,000, *d* = 1,000, and TV-distance 0.046, there are nearly twice as many singletons as under the Kingman (see (Figure 5d)). When *k* and *d* differ by nearly two orders of magnitude (*k* = 100, *d* = 9,900, TV-distance *<* 0.01; Figure 5a), the SFS under our model is close to the Kingman, although we still observe a small difference of roughly 0.15 times the number of singletons.

**Figure 5:**
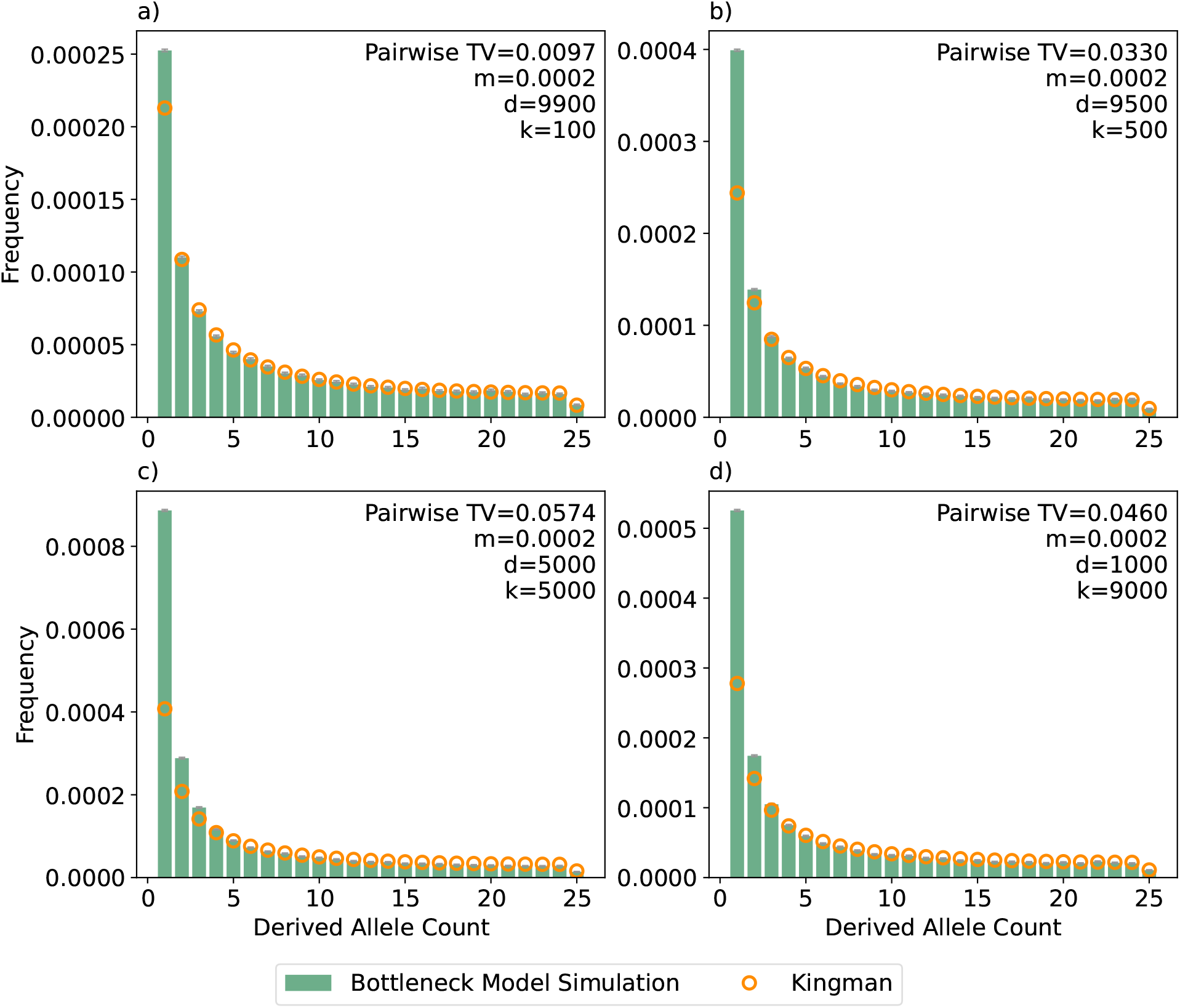
The simulated site frequency spectrum under our model calculated from 10,000 replicates (green bars) against the expected Kingman *θ*/*i* SFS with *θ* = *π_T_* (orange dots). The bottleneck size *k*, number of hosts *d*, and migration rate *m* are: (a) *k* = 100, *d* = 9,900, *m* = 0.0002; (b) *k* = 500, *d* = 9,500, *m* = 0.0002; (c) *k* = 5,000, *d* = 5,000, *m* = 0.0002; (d) *k* = 9,000, *d* = 1,000, *m* = 0.0002. In all panels, the mutation rate *u* = 10^−8^ and the host carrying capacity *N* = 10^10^. The pairwise TV-distance shown is calculated for each *k*, *d*, *m* combination.

We examine the relationship between the migration rate *m* and the deviation of the SFS under our model from the Kingman (Figure S3) by fixing *d* = *k* = 5,000 and varying *m*. The SFS deviates substantially from the Kingman when *m* = 0 (there are triple the singletons in our model as there are in the Kingman) and that we recover the Kingman when *m* ∼ 0.002 (2*dkm* = 10(*k* + *d*)) and TV-distance ∼ 0.003.

## Discussion

Because many pathogens experience recurrent bottlenecks while invading hosts, we used a coalescent framework to understand how such complex life cycles might affect patterns of neutral genetic diversity in pathogens. We employed a metapopulation model where each deme represents a host and migration between demes represents coinfection. We introduce two new assumptions in our model that differentiates ours from that of a standard island model. (1) We assume that all demes are equally likely to transmit their infection from one generation to another, thus allowing for stochasticity in pathogen transmission from one host to another. We refer to this as deme drift. (2) We assume that all demes undergo recurrent bottlenecks from *N* to *k* individuals every generation. This represents a transmission bottleneck during host invasion. We find that recurrent bottlenecks drastically reduce within-host diversity to the level of the bottleneck (*k*) when migration is low.

Under weak migration and when the size of the bottleneck (*k*) is of similar order to the number of infected hosts (*d*) in the population, the distribution of times to coalescence in the metapopulation diverges from that of a standard Wright-Fisher model and the site frequency spectrum is strongly skewed towards low frequency alleles (singletons). At the most extreme, we predict ∼3 times the singletons in our model as we would expect under the Kingman. This conflicts with some of the results of Gordo et al. (2009), who employed an SIR model to predict genetic diversity at equilibrium in a structured population. Under neutrality, they found values of Tajima’s *D* close to zero, indicating that the site frequency spectrum behaves similarly to the Kingman coalescent, and that this behavior does not depend on the effective size of a deme. However, they did not account for transmission bottlenecks, which might explain the difference in their observation.

### The effects of deme drift and recurrent bottlenecks

Recurrent bottlenecks reduce the level of within-host diversity by increasing the probability that pairs of lineages in the same deme coalesce. In the large-*N* limit, the probability that pairs of lineages in the same deme coalesce is approximately 1/*k*. Recurrent bottlenecks also reduce the amount of diversity in the whole metapopulation (*T_T_*) such that E [*T_T_*] depends on *k* + *d*. These approximations hold when 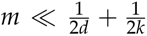. When *m* is large (panmixia) and *d*, *k* ≫ 1, E [*T_T_*] depends on the product 2*dkm*, and recurrent bottlenecks still reduce the level of diversity in the population. Recurrent bottlenecks also affect *F_ST_*: nucleotide diversity within a deme depends on *k* while nucleotide diversity in the whole population depends on *k* + *d* (assuming small *m*). When *k* ≪ *d*, *F_ST_* can be close to 1 even though the carrying capacity *N* of a deme is large.

Our results agree with Sigal et al. (2018), who use ordinary differential equations to model changes in the parasite population size within a host to predict levels of genetic diversity, that within-host diversity scales linearly with transmission bottleneck size. When the bottleneck size is one, every transmission bottleneck results in fixation and any diversity observed within a host is a result of *de novo* mutations. When the bottleneck size is greater than one, genetic diversity can be maintained within a host for more than one generation.

Deme drift, which allows parent demes to be chosen with replacement each generation, changes the time scale of the collecting process to be of order *d*. Specifically, the probability that two lineages in different demes enter the same deme in a single generation is 1/*d*. Note that this process is independent of the migration rate under our model; therefore *m* only affects the probability that two lineages in the same deme are placed into separate demes. This decoupling of the collecting process and the separating process allows the collecting process to occur faster than migration (which is of order 1/*m*) when the migration rate is small, resulting in coalescence to occur faster under our model.

### Convergence to the Kingman coalescent under some limits

Our model converges to the Kingman coalescent under two limits. These are: the limit where *N* = *k* → ∞ and *m* = 0 (the large deme limit; Notohara 1990) and the limit where *d* → ∞, *d* ≫ *k*, for any *m* (the many demes limit; Wakeley 1998). The second limit can be derived by following a separation of timescales argument where the collection probability of two states of lineages depends on what happens slowly at a timescale determined by *d* and the coalescence probability once two lineages collect happens quickly at timescale *k*. This can be compared to Wakeley’s many-demes limit except that the collecting probability does not depend on *m*, so *m* can be equal to 0. There is also the degenerate case where *d* = 1 and *N*, *k* → ∞.

### Population-genetic inference and confounding

A key observation from our model is that when the number of hosts is of similar magnitude to the number of pathogens during the bottleneck, the site frequency spectrum of neutral alleles is substantially skewed towards singletons. There are two implications of this. First, because the SFS at neutral sites is often used to infer ancestral population size changes (e.g., Gutenkunst et al. 2009; Excoffier et al. 2013), the standard population-genetic demographic inference approaches will be systematically biased and likely to result in inference of population expansion (which also leads to excessive singletons). Second, another process that leads to a large number of singletons is progeny skew (Eldon and Wakeley 2006; Birkner et al. 2013; Blath et al. 2016) i.e., a skewed offspring distribution, observed commonly in natural populations of many species like plants and viruses (Tellier and Lemaire 2014). Although skewed offspring distributions also result in a higher proportion of high-frequency derived alleles (Árnason et al. 2023), those are much harder to sample and characterize in population genetics data. Thus, a large excess of singletons can also be interpreted as a signature of strong progeny skew in the population. Our work here suggests that an SFS skewed toward low-frequency alleles can be generated in a simple metapopulation model where offspring distribution follows the Wright-Fisher model. Finally, a large proportion of singletons may also be caused by pervasive recurrent positive selection (Kim 2006), although we suggest caution against such an interpretation in pathogen populations where researchers are more likely to make such assumptions.

### Caveats to our model and future work

We discuss the following caveats and assumptions of our model that may inform future work.

#### Modeling recurrent bottlenecks

We assume that individuals in demes expand from size *k* to *N* in a single generation. In reality, pathogen populations in hosts grow exponentially or logistically. If we assume that pathogens only undergo migration (coinfection) at the beginning of an infection cycle, our results can be generalized to account for multiple replications in each infection cycle. Let pathogens replicate *j* times each generation, and let demes be of sizes *S*_1_, . . ., *S_j_* after each replication. During replications 1 to *j* − 1, demes are in a hidden state similar to the bottleneck in our model. Then, the probability that two lineages coalesce in a hidden state in a single generation is 1 − (1 − 1/*S*_1_) (1 − 1/*S*_2_) · · · 1 − 1/*S_j_*_−1_ and the probability that two lineages coalesce during replication *j* is 1/*S_j_*. These map directly onto 1/*k* and 1/*N* in our model by letting 1/*k* = 1 − (1 − 1/*S*_1_) (1 − 1/*S*_2_) · · · 1 − 1/*S_j_*_−1_ and *N* = *S_j_*. However, our results do not describe what happens when migration can occur at any time during the infection cycle.

#### Allowing for multiple mergers in pathogen transmission

For the purpose of this study, we have modeled deme drift such that all hosts are equally likely to transmit their infection to the next host, and that the number of individuals a host spreads their infection to is multinomially distributed. However, transmission in many pathogen populations may be highly skewed towards rare events where an infected host happens to infect many others by chance. This assumption of small progeny skew neglects superspreader events, which are known to occur in pathogens (Woolhouse et al. 1997; Stein 2011). Such transmission dynamics can easily be modeled in our framework by sampling from a skewed offspring distribution when incorporating deme drift. If transmission were highly skewed, we may expect to see multiple mergers even in small sample sizes. This may instead lead to the metapopulation SFS converging to that generated by a non-Kingman coalescent. It has been demonstrated that the Kingman coalescent is a poor choice of null model in the study of *Mycobacterium tuberculosis* due to multiple mergers (Menardo et al. 2021). Thus, future work in this area will allow for incorporating more realistic biological complexities of pathogen transmission in our model.

#### Overlapping generations and the Moran model

One direction of interest for future work is the generalization of this model to allow for overlapping generations. In an extreme example, if hosts transmit their infections such that each generation, one deme is removed from the metapopulation and is colonized by one of the other demes (with a bottleneck), host dynamics could take the form of a Moran model (Moran 1958). Hosts being removed from the population would indicate host death or the host being cured. More abstractly, it may be possible to describe an umbrella class of metapopulation models that undergo repeated generational bottlenecks; this model would contain the model we describe here as well as those with overlapping generations. In this example, it could be useful to model within-host pathogen dynamics according to a Wright-Fisher model and between-host dynamics according to a Moran model. Such a formulation would allow for overlapping generations to be considered, which can be especially relevant for pathogens that result in chronic infections.

#### Selection

We have only considered the dynamics of neutral mutations when modeling pathogen evolution. However, the dynamics of selected mutations, especially beneficial ones such as drug-resistance mutations, are of particular interest in pathogens. Roze et al. (2005) employed a diffusion framework to understand the properties of selected mutations in the same model as ours (including both deme drift and recurrent bottlenecks) when attempting to model mitochondrial populations. Their results suggest that beneficial mutations are less likely and mildly deleterious mutations are more likely to fix due to recurrent bottlenecks in comparison to a model without bottlenecks (also see Wahl and Gerrish 2001). In contrast, mathematical models of mitochondrial evolution that incorporate transmission bottlenecks (but not deme drift) suggest that selection against deleterious mutations may be more effective as the bottleneck size decreases due to greater variance in host fitness (Bergstrom and Pritchard 1998). It is unclear whether the times to fixation and loss of selected mutations are similarly affected by recurrent bottlenecks, which would play a role in genomic signatures generated by selective sweeps in pathogen populations. Future work in this area would be useful.

#### Allowing for uncertainty in flxed constants

Finally, we have made simplifying assumptions about the population being at equilibrium, with a fixed number of infected hosts in the metapopulation, while pathogen populations often have more complex demographic history (Altizer et al. 2006). Similarly, we assume a single constant size of the transmission bottlenecks that are fixed across all hosts. Modeling the bottleneck size as a distribution and allowing for non-equilibrium scenarios can be further extensions of our model for the future.

In summary, we provide a formal study of neutral diversity in pathogens using the coalescent. Further extensions of this work that include overlapping generations, progeny skew, and complex transmission dynamics will contribute to a more appropriate evolutionary null model for pathogen evolution. This will in turn benefit our understanding of the effects of selection on patterns of pathogen sequence variation data and allow for accurate population genomic inference.

## Methods

### Coalescent simulations of more than two samples

The expected site frequency spectrum (SFS) under our model is obtained using coalescent simulations of a sample size of *n >* 2. Our simulation tracks lineages as they move through a fixed number of demes, allowing for multiple coalescent events in a single generation and for coalescent events of more than two lineages. To initialize the simulation, each lineage is given a random deme with replacement. Every generation until all lineages have coalesced: Pairs of individuals in the same deme coalesce with probability 1/*k*. If a lineage is part of overlapping coalescent events, all coalescent events that lineage is involved in coalesce to a single lineage. Each lineage has probability *m* of being a migrant in any generation. Consistent with our model, all migrants in the same deme have the same parent deme. Demes independently sample a parent deme from the set {1, …, *d*} with replacement. Pairs of individuals in the same deme coalesce with probability 1/*N*. If a lineage is part of overlapping coalescent events, all coalescent events that lineage is involved in coalesce to a single lineage. The simulation is complete when all lineages have coalesced to a single ancestor.

Coalescent events add at least one edge to an output graph. This graph records the tree we simulate. The number of samples in each lineage is also recorded in the graph for calculation of the SFS. The SFS is calculated from the graph by summing the length of the edges with *i* descendants. Each bin of the site frequency spectrum, *ξ_i_*, is binomially distributed with probability *u* and *τ_i_* trials, where *τ_i_*represents the sum of the branch lengths with *i* descendants. Using the laws of total expectation and variance, we obtain

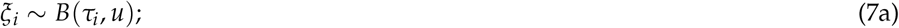

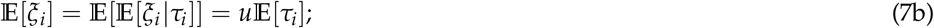

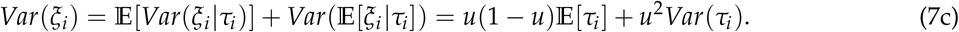

## Data Availability

All scripts to perform simulations are provided in the repository https://github.com/JohriLab/BottleneckModel.

## Acknowledgments

Research reported in this publication was supported by the National Institute of General Medical Sciences of the National Institutes of Health under award number R35GM154969 to PJ and in part by grants from the NSF (DMS-2235451) and Simons Foundation (MPS-NITMB-00005320) to the NSF-Simons National Institute for Theory and Mathematics in Biology (NITMB) (AM is affiliate faculty). The research in this study was conducted using computational resources provided by ITS Research Computing at the University of North Carolina at Chapel Hill. ChatGPT 4.0 and Claude 3.5 assisted with writing code and checking for mathematical errors.

## Notes

https://github.com/JohriLab/BottleneckModel

### Competing Interest Statement

The authors have declared no competing interest.

## References

1. Altizer, Sonia et al. 2006. Seasonality and the dynamics of infectious diseases. Ecol. Lett. 9(4): 467–484 10.1111/j.1461-0248.2005.00879.x.

2. Árnason, Einar, Koskela, Jere, Halldórsdóttir, Katrín and Eldon, Bjarki. 2023. Sweepstakes reproductive success via pervasive and recurrent selective sweeps. eLife. 12(e80781) 10.7554/eLife.80781.

3. Bergstrom, Carl T and Pritchard, Jonathan. 1998. Germline bottlenecks and the evolutionary maintenance of mitochondrial genomes. Genetics. 149(4): 2135–2146 10.1093/genetics/149.4.2135.

4. Birkner, Matthias, Blath, Jochen and Eldon, Bjarki. 2013. Statistical properties of the site-frequency spectrum associated with Λ-coalescents. Genetics. 195(3): 1037–1053 10.1534/genetics.113.156612.

5. Blath, Jochen, Cronjäger, Mathias Christensen, Eldon, Bjarki and Hammer, Matthias. 2016. The site-frequency spectrum associated with Ξ-coalescents. Theor. Popul. Biol. 110: 36–50 10.1016/j.tpb.2016.04.002.

6. Cao, Liqin et al. 2007. The mitochondrial bottleneck occurs without reduction of mtDNA content in female mouse germ cells. Nat. Genet. 39(3): 386–390 10.1038/ng1970.

7. Craven, Lyndsey et al. 2010. Pronuclear transfer in human embryos to prevent transmission of mitochondrial DNA disease. Nature. 465(7294): 82–85 10.1038/nature08958.

8. Eldon, Bjarki and Wakeley, John. 2006. Coalescent processes when the distribution of offspring number among individuals is highly skewed. Genetics. 172(4): 2621–2633 10.1534/genetics.105.052175.

9. Excoffier, Laurent, Dupanloup, Isabelle, Huerta-Sańchez, Emilia, Sousa, Vitor C. and Foll, Matthieu. 2013. Robust demographic inference from genomic and SNP data. PLOS Genet. 9(10): e1003905 10.1371/journal.pgen.1003905.

10. Floros, Vasileios I. et al. 2018. Segregation of mitochondrial DNA heteroplasmy through a developmental genetic bottleneck in human embryos. Nat. Cell Biol. 20(2): 144–151 10.1038/s41556-017-0017-8.

11. Fountain, Toby, Duvaux, Ludovic, Horsburgh, Gavin, Reinhardt, Klaus and Butlin, Roger K. 2014. Human-facilitated metapopulation dynamics in an emerging pest species, *Cimex lectularius*. Mol. Ecol. 23(5): 1071–1084 10.1111/mec.12673.

12. Fu, Y. X. 1995. Statistical properties of segregating sites. Theor. Popul. Biol. 48(2): 172–197 10.1006/tpbi.1995.1025.

13. Funk, Daniel J, Wernegreen, Jennifer J and Moran, Nancy A. 2001. Intraspecific variation in symbiont genomes: bottlenecks and the aphid-*Buchnera* association. Genetics. 157(2): 477–489 10.1093/genetics/157.2.477.

14. Ghafari, Mahan, Lumby, Casper K., Weissman, Daniel B. and Illingworth, Christopher J. R. 2020. Inferring transmission bottleneck size from viral sequence data using a novel haplotype reconstruction method. J. Virol. 94(13): 10.1128/jvi.00014–20 10.1128/jvi.00014-20.

15. Gordo, Isabel, Gomes, M. Gabriela M., Reis, Daniel G. and Campos, Paulo R. A. 2009. Genetic diversity in the SIR model of pathogen evolution. PLOS ONE. 4(3): e4876 10.1371/journal.pone.0004876.

16. Graumans, Wouter, Jacobs, Ella, Bousema, Teun and Sinnis, Photini. 2020. When is a *Plasmodium*-infected mosquito an infectious mosquito? Trends Parasitol. 36(8): 705–716 10.1016/j.pt.2020.05.011.

17. Griffiths, R. C. 2003. The frequency spectrum of a mutation, and its age, in a general diffusion model. Theor. Popul. Biol. 64(2): 241–251 10.1016/S0040-5809(03)00075-3.

18. Gutenkunst, Ryan N., Hernandez, Ryan D., Williamson, Scott H. and Bustamante, Carlos D. 2009. Inferring the joint demographic history of multiple populations from multidimensional SNP frequency data. PLOS Genet. 5(10): e1000695 10.1371/journal.pgen.1000695.

19. Henry, Cobi M. et al. 2025 Dec 22. Towards an evolutionary baseline model of *Plasmodium falciparum* for population-genomic inference. 10.64898/2025.12.20.695730.

20. Kappe, Stefan H. I., Vaughan, Ashley M., Boddey, Justin A. and Cowman, Alan F. 2010. That was then but this is now: malaria research in the time of an eradication agenda. Science. 328(5980): 862–866.

21. Kennedy, David A. and Dwyer, Greg. 2018. Effects of multiple sources of genetic drift on pathogen variation within hosts. PLOS Biol. 16(3): e2004444 10.1371/journal.pbio.2004444.

22. Kim, Yuseob. 2006. Allele frequency distribution under recurrent selective sweeps. Genetics. 172(3): 1967–1978 10.1534/genetics.105.048447.

23. Kingman, J. F. C. 1982. On the genealogy of large populations. J. Appl. Probab. 19: 27–43 10.2307/3213548.

24. Latter, B. D. H. 1973. The island model of population differentiation: a general solution. Genetics. 73(1): 147–157 10.1093/genetics/73.1.147.

25. Lythgoe, Katrina A. et al. 2021. SARS-CoV-2 within-host diversity and transmission. Science. 372(6539): eabg0821 10.1126/science.abg0821.

26. Malaria Genomic Epidemiology Network, (MalariaGEN) et al. 2025. Pf8: an open dataset of *Plasmodium falciparum* genome variation in 33,325 worldwide samples. Wellcome Open Res. 10: 325 10.12688/wellcomeopenres.24031.1.

27. Maruyama, Takeo. 1970. Effective number of alleles in a subdivided population. Theor. Popul Biol. 1(3): 273–306 10.1016/0040-5809(70)90047-X.

28. McCrone, John T et al. 2018. Stochastic processes constrain the within and between host evolution of influenza virus. eLife. 7: e35962 10.7554/eLife.35962.

29. Menardo, Fabrizio, Gagneux, Sébastien and Freund, Fabian. 2021. Multiple merger genealogies in outbreaks of *Mycobacterium tuberculosis*. Mol. Biol. Evol. 38(1): 290–306 10.1093/molbev/msaa179.

30. Moran, P. A. P. 1958. Random processes in genetics. Math. Proc. Cambridge Philos. Soc. 54(1): 60–71 10.1017/S0305004100033193.

31. Moran, P. A. P. 1959. The theory of some genetical effects of population subdivision. Aust. J. Biol. Sci. 12(2): 109–116 10.1071/BI9590109.

32. Nagylaki, Thomas. 1998. The expected number of heterozygous sites in a subdivided population. Genetics. 149(3): 1599–1604 10.1093/genetics/149.3.1599.

33. Notohara, M. 1990. The coalescent and the genealogical process in geographically structured population. J. Math. Biol. 29(1): 59–75 10.1007/BF00173909.

34. Pannell, John R. 2003. Coalescence in a metapopulation with recurrent local extinction and recolonization. Evolution. 57(5): 949–961 10.1111/j.0014-3820.2003.tb00307.x.

35. Pietri, Jose E., DeBruhl, Heather and Sullivan, William. 2016. The rich somatic life of *Wolbachia*. MicrobiologyOpen. 5(6): 923–936 10.1002/mbo3.390.

36. Roze, Denis, Rousset, Francois and Michalakis, Yannis. 2005. Germline bottlenecks, biparental inheritance and selection on mitochondrial variants. Genetics. 170(3): 1385–1399 10.1534/genetics.104.039495.

37. Sender, Ron et al. 2021. The total number and mass of SARS-CoV-2 virions. Proc. Natl. Acad. Sci. U.S.A. 118(25): e2024815118 10.1073/pnas.2024815118.

38. Sigal, Daniel, Reid, Jennifer N S and Wahl, Lindi M. 2018. Effects of transmission bottlenecks on the diversity of influenza A virus. Genetics. 210(3): 1075–1088 10.1534/genetics.118.301510.

39. Smith, Emily A., Fleming, Derek F., Lackritz, Eve M. and Ulrich, Angela K. 2026. Inequities and global declines in SARS-CoV-2 genomic data availability hinder response to emerging variants. Npj Viruses. 4(1): 13 10.1038/s44298-026-00176-7.

40. Stein, Richard A. 2011. Super-spreaders in infectious diseases. Int. J. Infect. Dis. 15(8): e510–e513 10.1016/j.ijid.2010.06.020.

41. Tajima, F. 1989. Statistical method for testing the neutral mutation hypothesis by DNA polymorphism. Genetics. 123(3): 585–595 10.1093/genetics/123.3.585.

42. Tellier, Aurélien and Lemaire, Christophe. 2014. Coalescence 2.0: a multiple branching of recent theoretical developments and their applications. Mol. Ecol. 23(11): 2637–2652 10.1111/mec.12755.

43. To, Kelvin K.W. et al. 2010. Viral load in patients infected with pandemic H1N1 2009 influenza A virus. J. Med. Virol. 82(1): 1–7 10.1002/jmv.21664.

44. Wahl, Lindi M. and Gerrish, Philip J. 2001. The probability that beneficial mutations are lost in populations with periodic bottlenecks. Evolution. 55(12): 2606–2610 10.1111/j.0014-3820.2001.tb00772.x.

45. Wakeley, John. 1998. Segregating sites in Wright’s island model. Theor. Popul. Biol. 53(2): 166–174 10.1006/tpbi.1997.1355.

46. Wakeley, John and Aliacar, Nicolas. 2001. Gene genealogies in a metapopulation. Genetics. 159(2): 893–905 10.1093/genetics/159.2.893.

47. Woolhouse, M. E. J. et al. 1997. Heterogeneities in the transmission of infectious agents: implications for the design of control programs. Proc. Natl. Acad. Sci. U.S.A. 94(1): 338–342 10.1073/pnas.94.1.338.

48. Wright, Sewall. 1931. Evolution in mendelian populations. Genetics. 16(2): 97–159 10.1093/genetics/16.2.97.

49. Zaidi, Arslan A. et al. 2019. Bottleneck and selection in the germline and maternal age influence transmission of mitochondrial DNA in human pedigrees. Proc. Natl. Acad. Sci. U.S.A. 116(50): 25172–25178 10.1073/pnas.1906331116.

50. Zhang, Haixin, Burr, Stephen P. and Chinnery, Patrick F. 2018. The mitochondrial DNA genetic bottleneck: inheritance and beyond. Essays Biochem. 62(3): 225–234 10.1042/EBC20170096.

