## Supplement for "A model for within- and between-host evolution in pathogens"

### Supplementary Figures

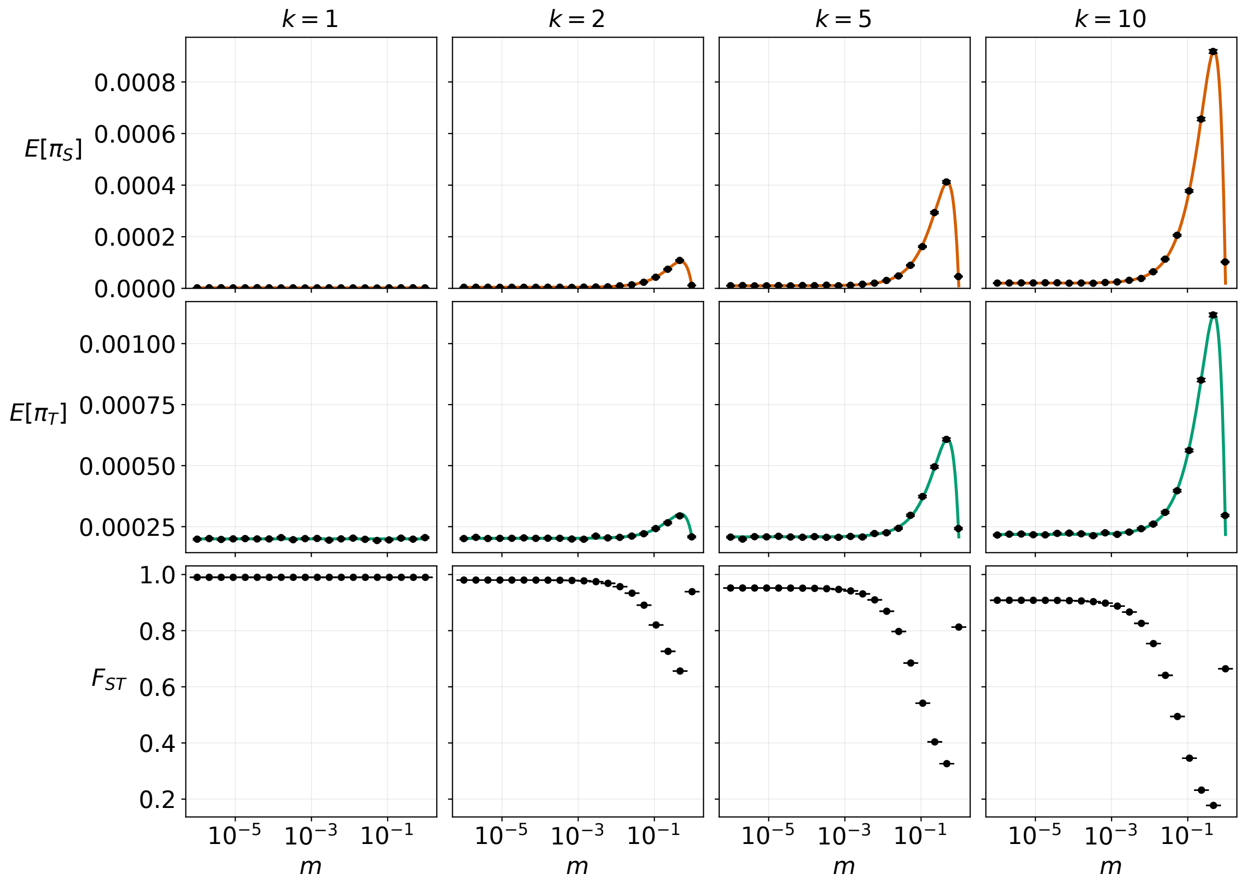


Figure S1: Comparison of theoretically calculated $\pi_S$ and $\pi_T$ with results from 20,000,000 ($= n_r$) replicates of coalescent simulations for varying bottleneck sizes ($k$) and migration rates ($m$). Here $d = 100$ and $N = 10^{10}$. The theoretical means of each statistic are given by the lines, while the ribbons represent the expectation of the standard error $\frac{\sigma}{\sqrt{n_r}}$. The simulated means are given by the points, while the error bars represent the standard error. As $n_r \to \infty$ the standard errors approach zero, and the simulated means converge to the theoretical expectations.

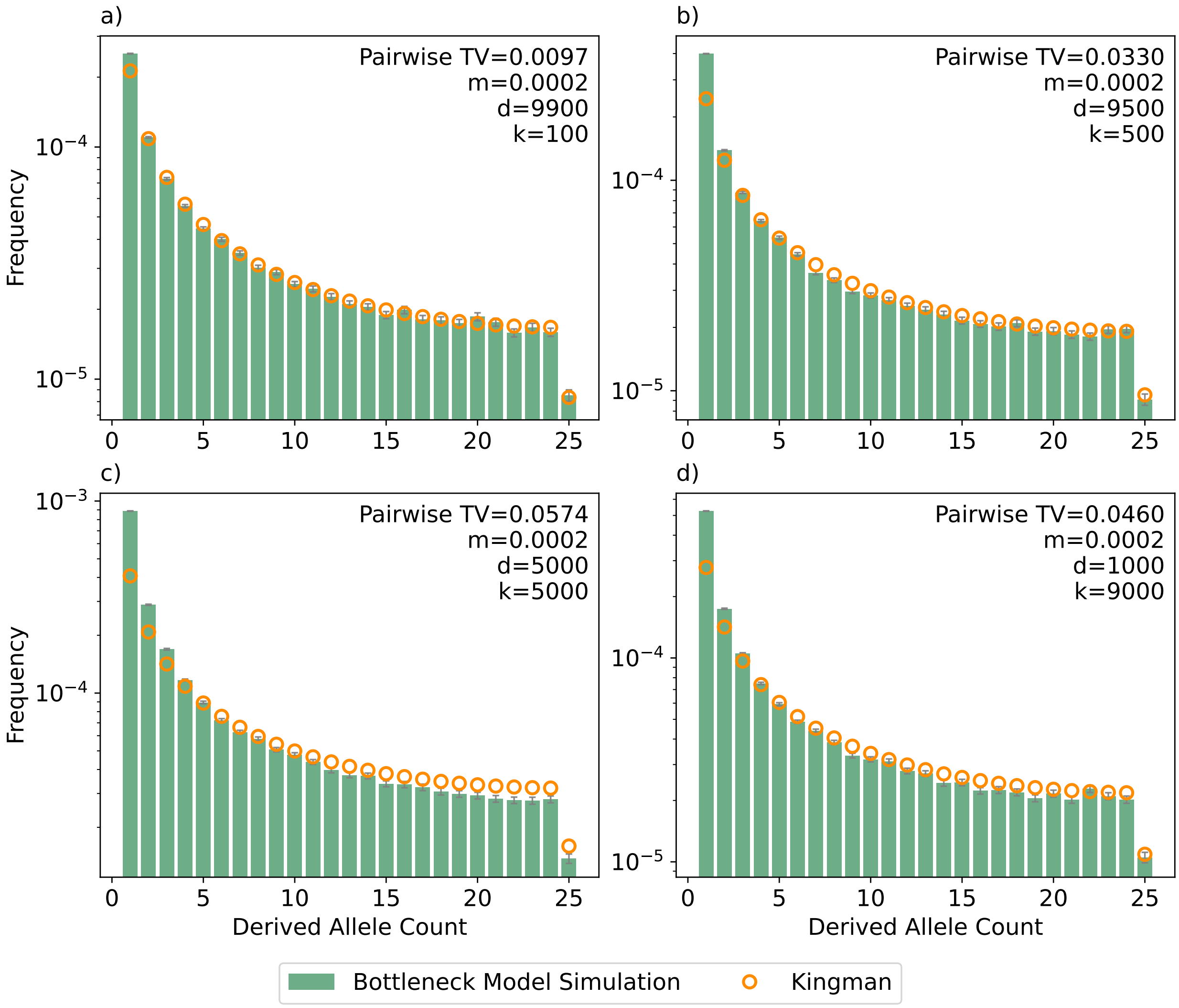


Figure S2: The simulated site frequency spectrum under our model calculated from 10,000 replicates (green bars) against the expected Kingman $\theta/i$ SFS with $\theta = \pi_T$ (orange dots). The y-axis is on a log-scale. The bottleneck size $k$, number of hosts $d$, and migration rate $m$ are: (a) $k = 100$, $d = 9900$, $m = 0.0002$; (b) $k = 500$, $d = 9500$, $m = 0.0002$; (c) $k = 5000$, $d = 5000$, $m = 0.0002$; (d) $k = 9000$, $d = 1000$, $m = 0.0002$. In all panels, the mutation rate $u = 10^{-8}$ and the host carrying capacity $N = 10^{10}$. The pairwise TV-distance shown is calculated for each $k, d, m$ combination.

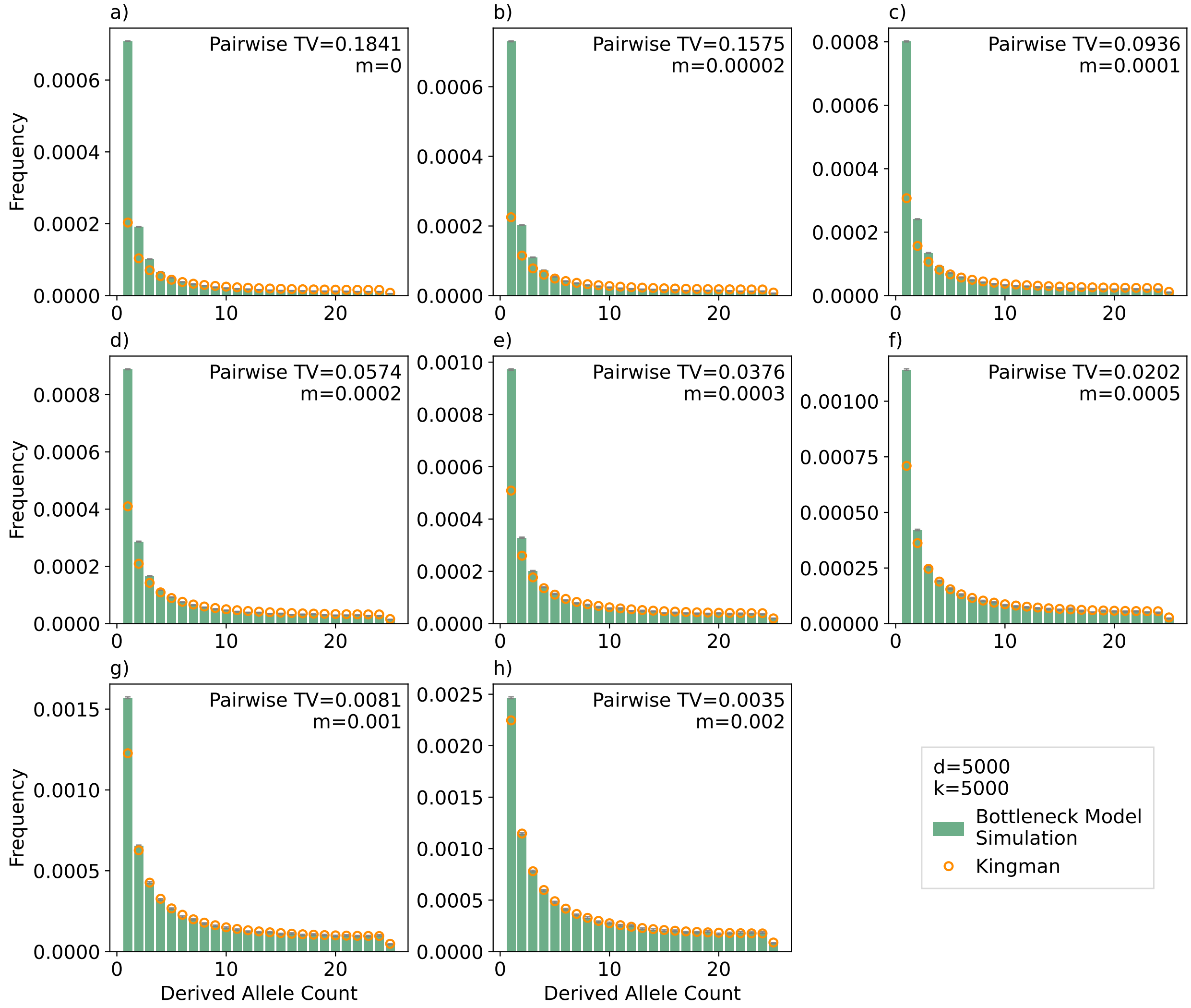


Figure S3: The simulated site frequency spectrum under our model calculated from 10,000 replicates (green bars) against the expected Kingman $\theta/i$ SFS with $\theta = \pi_T$ (orange dots). The bottleneck size $k$ and number of hosts $d$ are fixed at 5000, and the varying migration rates $m$ are shown in the top-right portion of each panel. In all panels, the mutation rate $u = 10^{-8}$ and the host carrying capacity $N = 10^{10}$. The pairwise TV-distance shown is calculated for each $k, d, m$ combination.

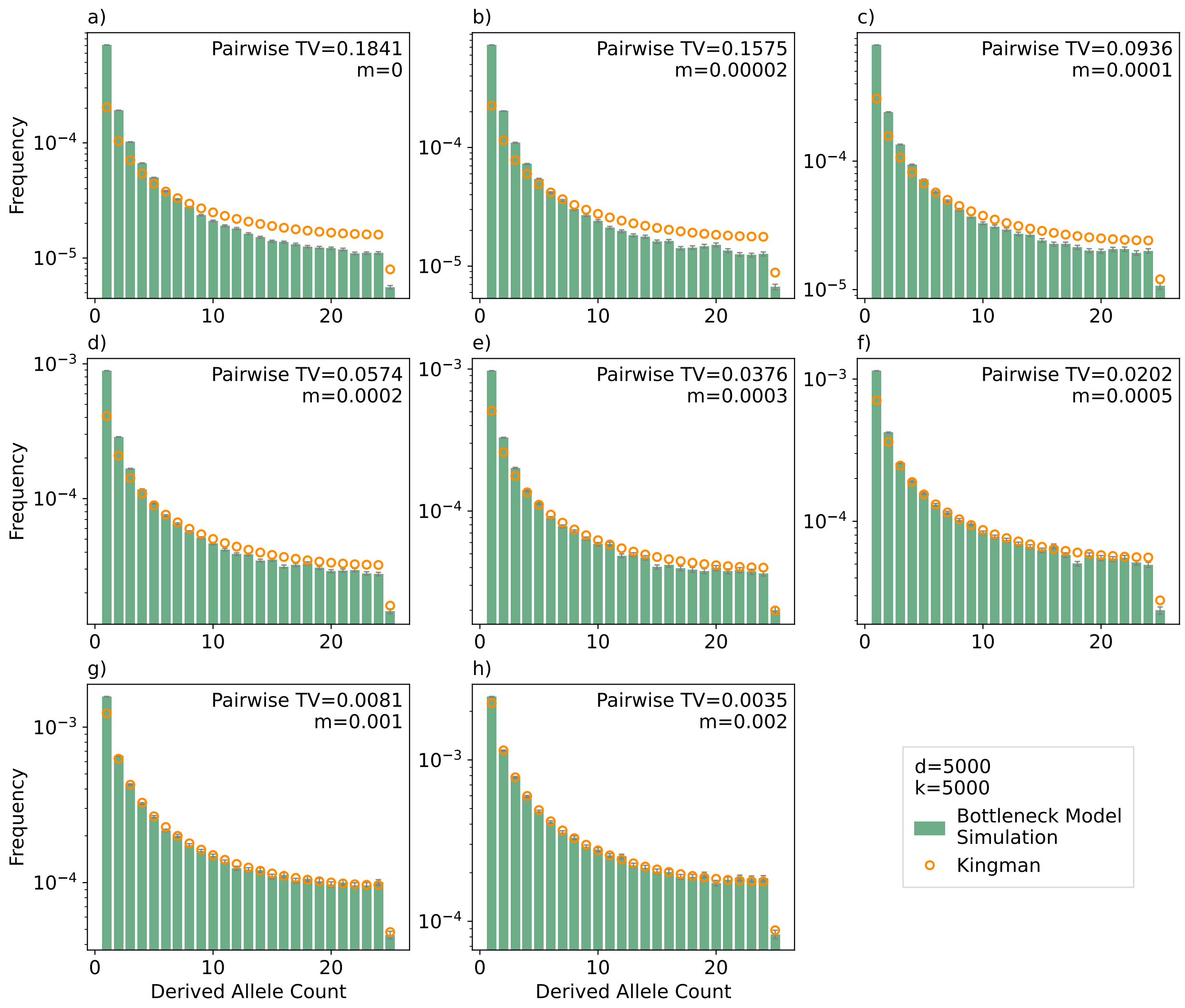


Figure S4: The simulated site frequency spectrum under our model calculated from 10,000 replicates (green bars) against the expected Kingman $\theta/i$ SFS with $\theta = \pi_T$ (orange dots). They y-axis is on a log-scale. The bottleneck size $k$ and number of hosts $d$ are fixed at 5000, and the varying migration rates $m$ are shown in the top-right portion of each panel. In all panels, the mutation rate $u = 10^{-8}$ and the host carrying capacity $N = 10^{10}$. The pairwise TV-distance shown is calculated for each $k, d, m$ combination.

### Supplementary Text

**Text S1: Coalescent simulations of a pair of samples.**

1. We choose whether the pair of lineages starts in the same deme or in different demes. To simulate $\pi_S$ the individuals begin in the same deme. To simulate $\pi_T$ the individuals begin in the same deme with probability $\frac{1}{d}$.

2. Every generation until the pair of lineages coalesces:

   - If the pair is in the same deme:

     (a) The pair coalesces immediately with probability $\frac{1}{k}$. If the pair coalesces, the simulation ends.

     (b) Exactly one lineage migrates out of the deme with probability $\alpha$. If one lineage is a migrant, the pair of lineages are now in different demes.

     (c) If the pair remain in the same deme, they coalesce immediately with probability $\frac{1}{N}$. If they coalesce, the simulation ends.

   - If the pair are in different demes:

     (a) The pair of lineages drift to the same deme with probability $\frac{1}{d}$.

     (b) If the pair of lineages is now in the same deme, they coalesce immediately with probability $\frac{1}{N}$. If they coalesce, the simulation ends.

   - The number of generations until the pair of lineages coalesces is recorded. The number of mutations between the two lineages ($\pi$) is drawn from a binomial distribution with probability $u$ and $2T$ trials.
